# Comparison of Urinary Proteome Modifications Between Methamphetamine-Abstinent Rehabilitation Patients for Over Three Months and Healthy Individuals

**DOI:** 10.1101/2025.04.02.646921

**Authors:** Yuzhen Chen, Ziyun Shen, Juncheng Liang, Yanping Deng, Youhe Gao

## Abstract

Methamphetamine addiction is characterized by complex mechanisms and a high relapse rate, posing a serious public health threat. In this study, urinary proteome modifications of rehabilitation patients who had abstained from methamphetamine for over three months and those of healthy individuals were analyzed and compared. A total of 984 differentially modified peptides were identified, 362 of which showed changes from presence to absence or vice versa. Results from the randomized grouping test indicated that at least 74.85% of these differentially modified peptides were not randomly generated. Several proteins containing these peptides have been reported to be associated with methamphetamine, including prostaglandin-H2 D-isomerase, complement factor H, filamin-A, and plasminogen. Additionally, some significantly changed modified peptides and the proteins they are located in, which have not been previously reported to be associated with methamphetamine, may provide new insights into addiction rehabilitation. This study establishes a method for investigating drug addiction through urinary proteome modifications and observes differences between methamphetamine-abstinent rehabilitation patients and healthy individuals, which may help explain the drug’s high relapse rate. It offers a new perspective on drug withdrawal research and holds potential for tracking and assessing the rehabilitation process of methamphetamine addiction.

## 1. Introduction

Methamphetamine, a highly addictive psychostimulant, poses a serious global public health threat. According to the 2024 World Drug Report, the number of amphetamine-type stimulant (ATS) users worldwide was estimated to reach 30 million in 2022. Methamphetamine dominates the ATS market, which continues to expand in the Near and Middle East and South-West Asia. Additionally, methamphetamine production in the Golden Triangle region of Southeast Asia has surged in recent years, surpassing traditional opiates such as heroin and opium [1].

Methamphetamine’s high lipophilicity allows it to rapidly cross the blood-brain barrier and enter the central nervous system [2]. It increases the level and activity of three major monoamines—dopamine, serotonin, and norepinephrine in the brain’s reward circuits [3]. These effects contribute to its strong addictive potential. Long-term methamphetamine abuse can lead to severe physical and mental health consequences, including an increased risk of psychosis, depression, suicidal tendencies [4], cardiovascular diseases [5], and sexually transmitted infections [6]. It significantly reduces users’ quality of life and can be life-threatening.

Methamphetamine addicts are highly susceptible to relapse even after undergoing treatment. A study of 350 methamphetamine users in Los Angeles County found that 36% of patients relapsed within the first month of treatment, 14% relapsed within 2-6 months, and 11% within 7-12 months, while only 13% maintained abstinence for at least 5 years [7]. These findings highlight the urgent need for new drug addiction prevention and treatment strategies.

Even after withdrawal, it is still difficult for methamphetamine-addicted patients to return to normal level within a short period. Studies have shown that while thalamic metabolic activity recovers after 14 months of withdrawal, metabolic activity in the striatum, especially in the caudate nucleus and nucleus accumbens, remains lower than in healthy controls [8]. Long-term methamphetamine use also results in hard-to-recover cortical thickening in bilateral superior frontal gyri, as well as recoverable volumetric reduction in right hippocampus, bilateral accumbens nuclei, and bilateral cortical regions around insulae [9]. Furthermore, endogenous metabolites in the plasma, serum, and urine of methamphetamine addicts, rehabilitation patients, and healthy individuals were analyzed. Partial least squares discriminant analysis demonstrated clear separation among the acute, rehabilitation, and healthy groups. Additionally, abnormalities were observed in fatty acid metabolism, sulfate/sulfite metabolism, and sex hormone metabolism in the rehabilitation group [10]. These findings suggest that differences still exist between methamphetamine rehabilitation patients and healthy individuals.

Proteomics investigates the composition and changing laws of proteins in cells or organisms by analyzing protein structure, expression, post-translational modifications, and interactions [11]. As a filtrate of blood, urine does not need or possess homeostatic mechanisms to maintain stability. Thus, urine can accommodate and accumulate more changes without harming the body, reflecting changes in all organs and systems of the body earlier and more sensitively. Therefore, urine is a better source of biomarkers [12]. Studies utilizing urinary proteome reflect changes in the body in a comprehensive and systematic way.

In our previous research, we explored urinary proteome changes in methamphetamine addicts and rehabilitation patients who had abstained from methamphetamine for over three months, identifying significant differences between the rehabilitation patients and healthy individuals. Some differential proteins and pathways in these rehabilitation patients overlapped with those in acute-phase drug users [13]. In this study, we explored the differences between rehabilitation patients who had abstained from methamphetamine for over three months and healthy individuals from the perspective of urinary proteome modifications for the first time, providing a new perspective on the study of drug withdrawal (Figure 1).

**Figure 1.**
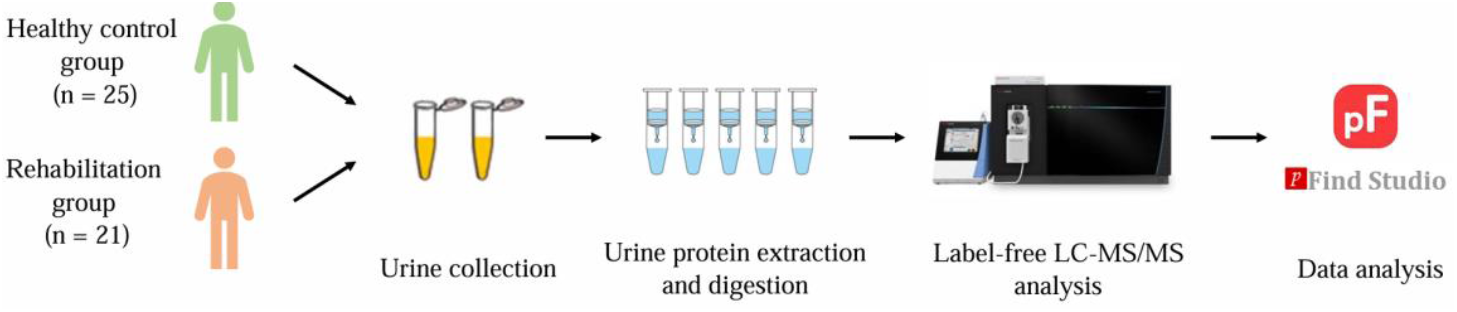
Technical workflow for comparing the urinary proteome modifications between methamphetamine-abstinent rehabilitation patients and healthy individuals.

## 2. Materials and Methods

### 2.1 Urine Sample Information

The mass spectrometry data used in this study were obtained from the research made publicly available online [13]. The control group consisted of healthy volunteers, while the rehabilitation group included drug rehabilitation patients who had abstained from methamphetamine for over three months. Samples with low peptide content or from patients with other diseases were excluded. Ultimately, 25 samples from the healthy control group and 21 samples from the rehabilitation group were included in the analysis. Sample information is provided in Table 1.

**Table 1.** Sample information of the rehabilitation and healthy control groups.

| Age | Healthy control group (n = 25) | Rehabilitation group (n = 21) |
| --- | --- | --- |
| Age range | 24~42 | 24~49 |
| Mean ± SD | 30.8±5.56 | 32.9±7.08 |

### 2.2 Database Searching and Data Processing

Modification information of the proteome was obtained using pFind Studio software (version 3.2.0, Institute of Computing Technology, Chinese Academy of Sciences, Beijing, China). Data collected by the Orbitrap Fusion Lumos Tribrid Mass Spectrometer were analyzed for label-free quantification, with the original data files searched against the *Homo sapiens* UniProt canonical database (updated in September 2024). For the search, “HCD-FTMS” was selected for “MS Instrument”, “Trypsin_P KR P C” was chosen for trypsin digestion, and the maximum number of missed cleavages allowed per peptide was two. Both precursor and fragment tolerances were set to ±20 ppm. To identify global modifications, “Open Search” was selected. False discovery rates (FDRs) at the spectra, peptide, and protein levels were all kept below 1%. The number of peptide mass spectra (Total_spec_num@pep) in each sample was extracted from the analysis results in pFind Studio using the script “pFind_protein_contrast_script.py” [14,15].

### 2.3 Data Analysis

The number of modified peptide mass spectra identified in the rehabilitation and healthy control groups were compared. Differentially modified peptides were screened with the following criteria: fold change (FC) ≥1.5 or ≤ 0.67, and *p* < 0.05 by two-tailed unpaired *t*-test analysis.

Hierarchical cluster analysis (HCA) and principal component analysis (PCA) were performed using the ‘Wu Kong’ platform (https://www.omicsolution.com/wkomics/main/). Biological analysis was conducted using the UniProt website (https://www.uniprot.org/). Functional analysis was supported by literature searches in the PubMed database (https://pubmed.ncbi.nlm.nih.gov).

## 3. Results and Discussion

### 3.1 Identification of Differentially Modified Peptides

Using a label-free quantitative proteome method, data from 46 samples were obtained by LC-MS/MS analysis. A search based on open-pFind yielded detailed information on the mass spectra number of modified peptides in each sample, including the proteins they were located in and the types of modifications contained in the peptides. A total of 45,057 modified peptides were identified, with 984 differentially modified peptides screened out based on the screening criteria of FC ≥ 1.5 or ≤ 0.67 and *p* < 0.05. The differentially modified peptides with significant changes are listed in Table 2. These peptides simultaneously meet the following conditions: an extremely large or small FC value, a very small *p*-value, and a large number of samples in at least one group (healthy control or rehabilitation group) in which the modified peptides could be identified. Detailed information of all differentially modified peptides is listed in Table S1, including peptide sequences, modification types, and the proteins containing these peptides.

**Table 2.**
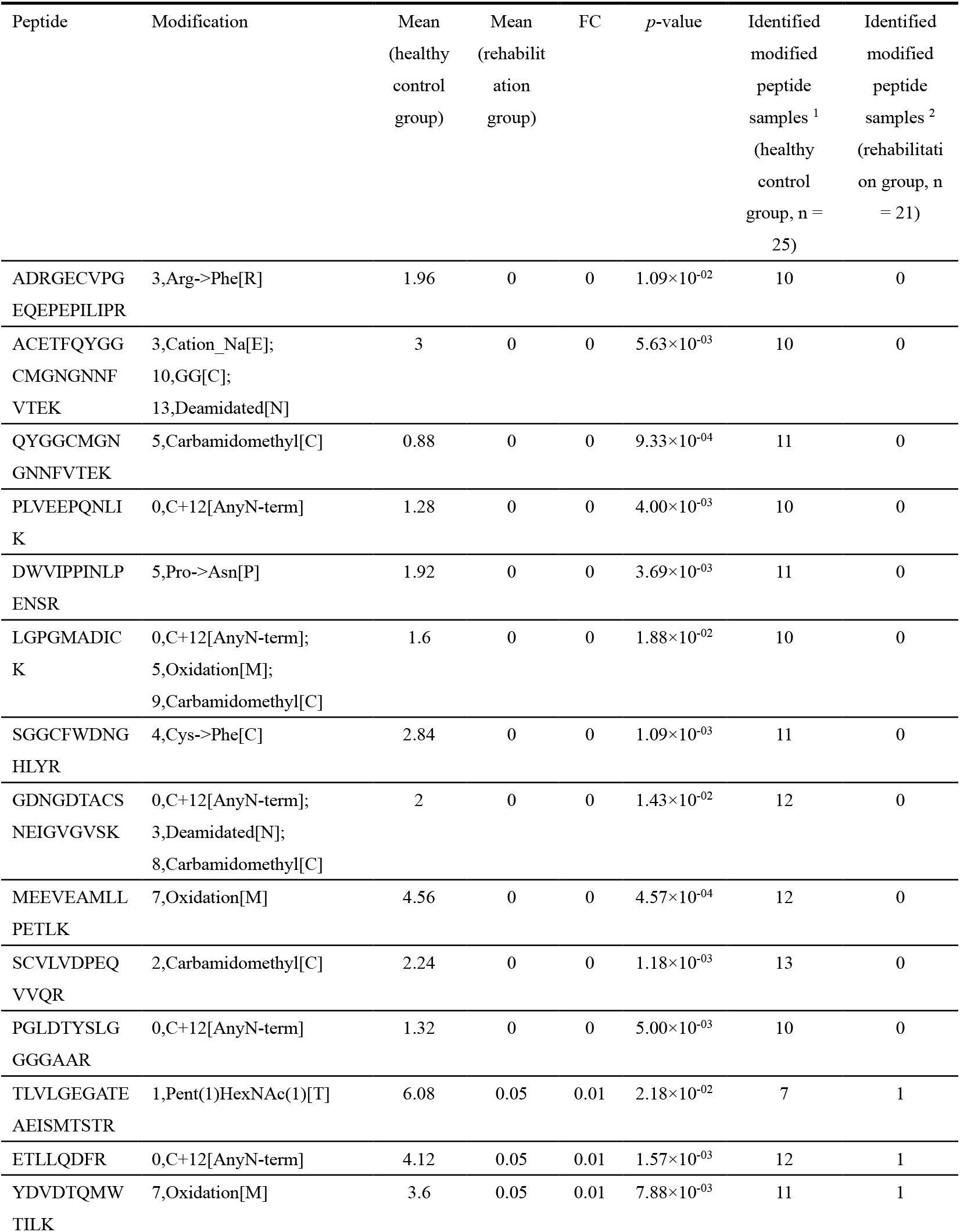
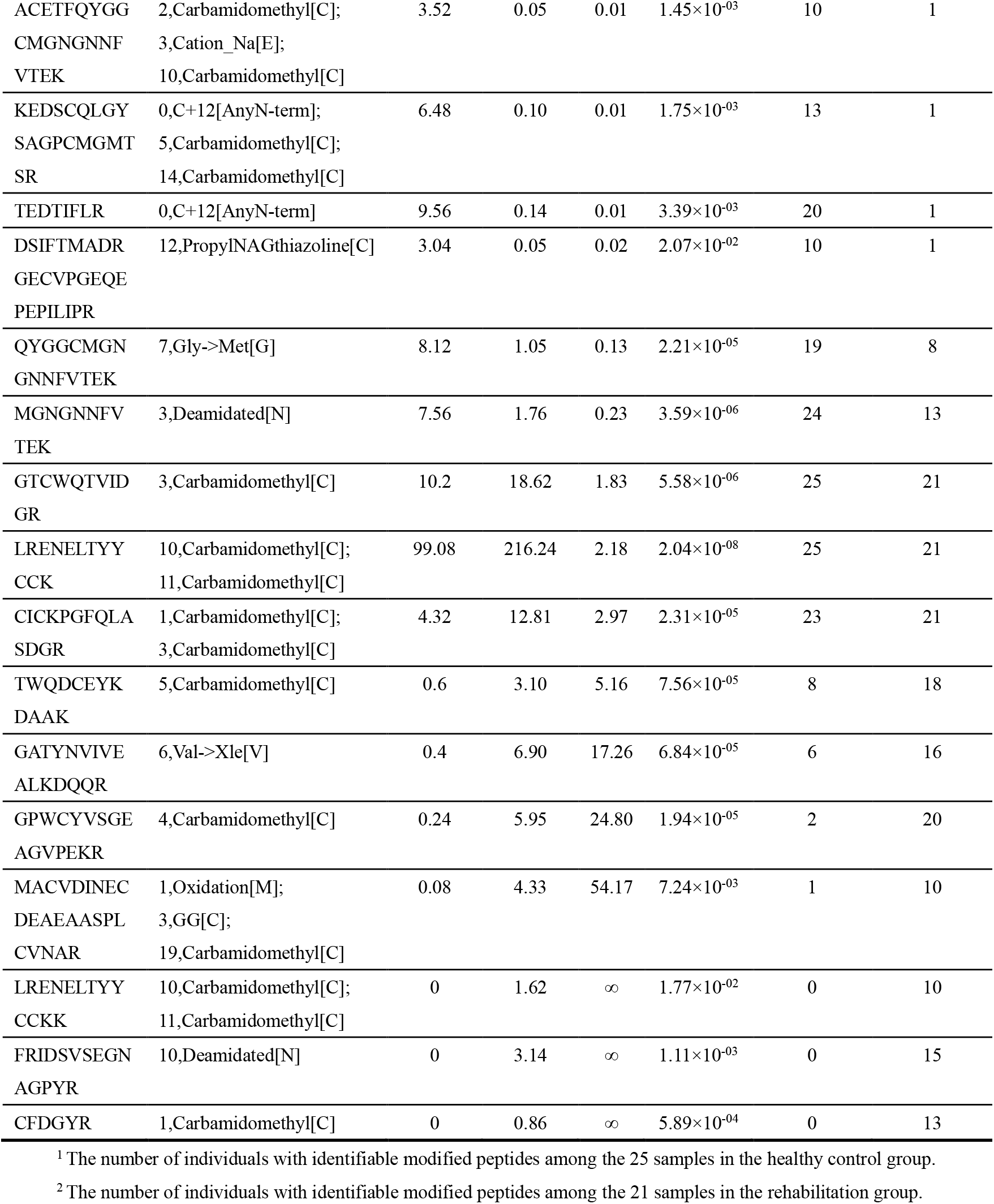
Differentially modified peptides with significant changes.

Among the 984 differentially modified peptides, 278 showed a change from presence to absence. This means that these peptides were identified in some of the 25 samples from the healthy control group but were not detected in any of the 21 samples from the rehabilitation group. Additionally, 84 peptides showed a change from absence to presence. Specifically, these peptides were identified in some of the 21 samples from the rehabilitation group but were not detected in any of the 25 samples from the healthy control group.

HCA and PCA were performed on the identified differentially modified peptides, and both of which distinguished the samples from the healthy control and rehabilitation groups (Figures 2 and 3).

**Figure 2.**
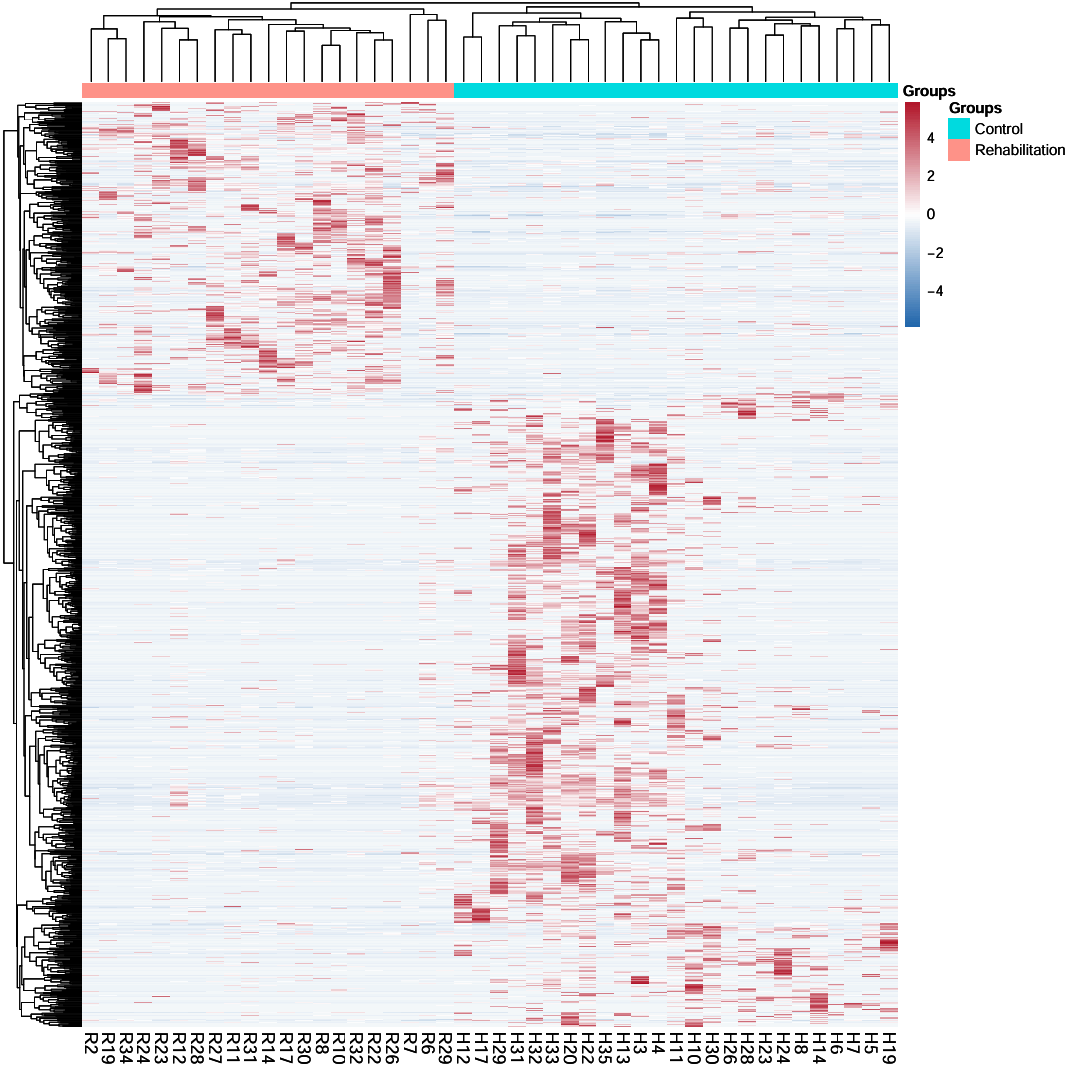
HCA of differentially modified peptides distinguished the samples from the healthy control and rehabilitation groups.

**Figure 3.**
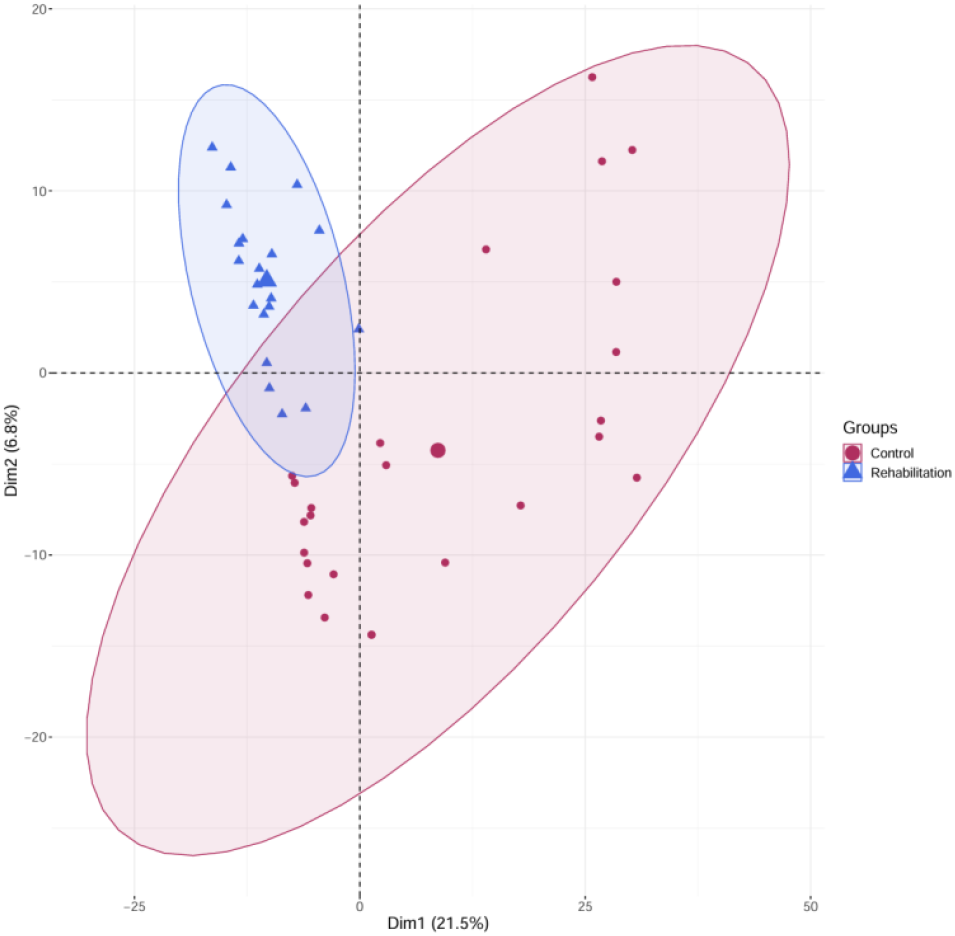
PCA of differentially modified peptides distinguished the samples from the healthy control and rehabilitation groups.

### 3.2. Randomized Grouping Test for Total Modified Peptides

To assess the possibility of random generation of the identified differentially modified peptides, a randomized grouping test was performed on the total modified peptides. Due to limitations in the computing power of the computers used in our laboratory, it was not feasible to perform the randomized grouping test on all samples. Therefore, we wrote code to randomly select 17 out of 25 samples from the healthy control group and 14 out of 21 samples from the rehabilitation group. This selection scale represents the maximum sample size that could support the randomized grouping test under the current conditions of the computing power in our laboratory. This approach ensures that, within the limitations of existing technologies, the study can still obtain reliable data to the greatest extent possible. The 31 selected samples were randomly divided into two new groups, resulting in a total of 265,182,525 combinations. These combinations were then screened for differences based on the same criteria (FC ≥ 1.5 or ≤ 0.67, *p* < 0.05). Similarly, limited by the computing power, we sampled the total combinations in intervals of 200,000, yielding 1,326 combinations. The number of randomly generated differentially modified peptides was then counted. The sampling process was repeated 10 times, and the randomized grouping test was performed after each sampling. Finally, the mean percentage of reliable differentially modified peptides was calculated. The results showed that at least 74.85% of the differentially modified peptides were not randomly generated, with a standard deviation of 6.95% (Table 3).

**Table 3.** Results of randomized grouping test.

| Sampling group | Number of differentially modified peptides | Number of differentially modified peptides generated randomly | Percentage of reliable differentially modified peptides |
| --- | --- | --- | --- |
| 1 | 383 | 126.88 | 66.87% |
| 2 | 450 | 143.25 | 68.17% |
| 3 | 623 | 140.70 | 77.42% |
| 4 | 516 | 145.08 | 71.88% |
| 5 | 1100 | 139.48 | 87.32% |
| 6 | 528 | 130.39 | 75.31% |
| 7 | 489 | 141.01 | 71.16% |
| 8 | 694 | 141.56 | 79.60% |
| 9 | 398 | 128.67 | 67.67% |
| 10 | 909 | 154.06 | 83.05% |

### 3.3. Analysis of Differential Modifications

The types and identification times of each modification in the 984 differentially modified peptides were counted. A total of 343 modification types were identified, with detailed information provided in Table S2. The modification types with ≥7 identifications are listed in Table 4.

**Table 4.**
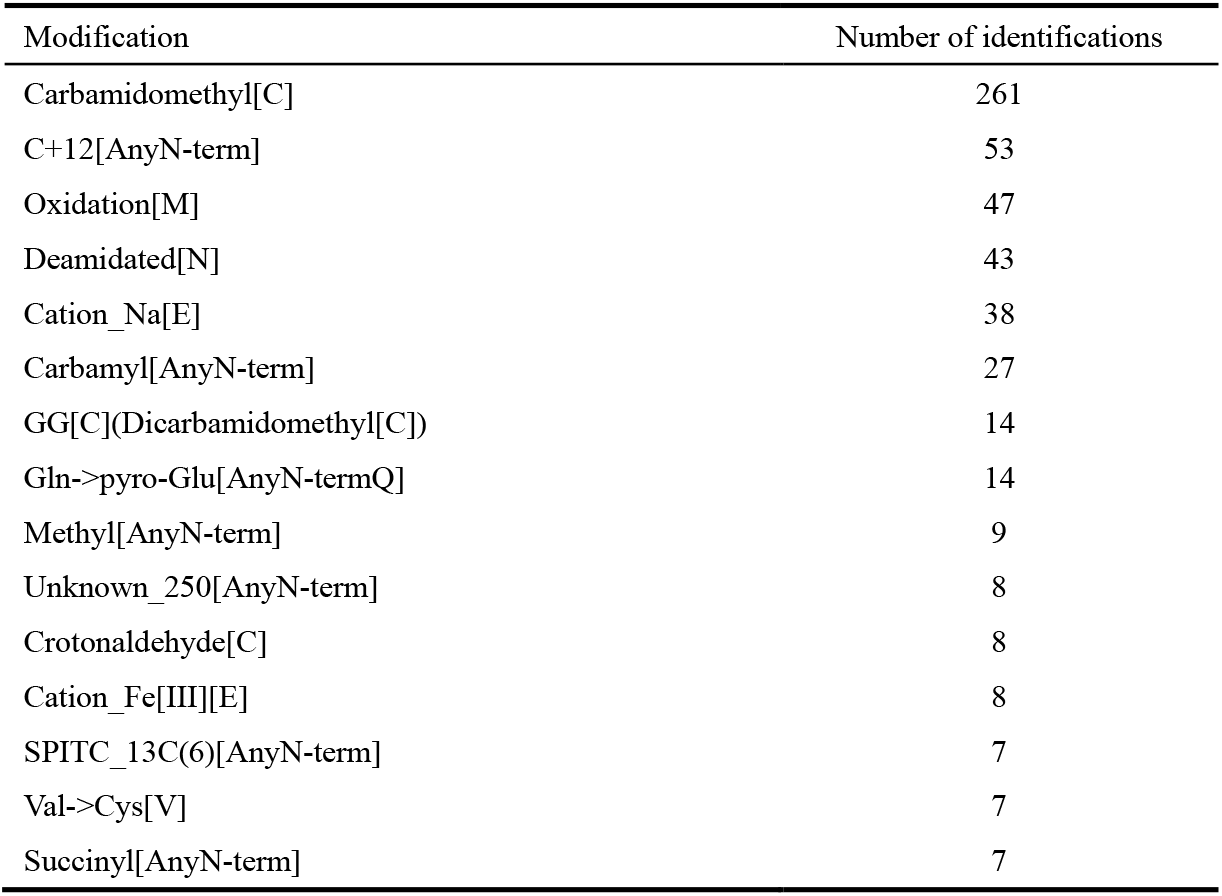
Differential modifications identified ≥ 7 times.

| Modification | Number of identifications |
| --- | --- |
| Carbamidomethyl[C] | 261 |
| C+12[AnyN-term] | 53 |
| Oxidation[M] | 47 |
| Deamidated[N] | 43 |
| Cation_Na[E] | 38 |
| Carbamyl[AnyN-term] | 27 |
| GG[C](Dicarbamidomethyl[C]) | 14 |
| Gln->pyro-Glu[AnyN-termQ] | 14 |
| Methyl[AnyN-term] | 9 |
| Unknown_250[AnyN-term] | 8 |
| Crotonaldehyde[C] | 8 |
| Cation_Fe[III][E] | 8 |
| SPITC_13C(6)[AnyN-term] | 7 |
| Val->Cys[V] | 7 |
| Succinyl[AnyN-term] | 7 |

The most frequent modification is carbamidomethyl, which is induced by the alkylation reagent iodoacetamide (IAA) acting on cysteine residues (Cys, C). This is an artificially introduced modification. In this study, samples from both the healthy control and rehabilitation groups were treated in the same manner. If the state of the cysteine residues in both groups had been identical prior to treatment, no differences should have been observed. However, the results showed that this modification was identified as different between the two groups of samples an extremely high number of times. This suggests a pre-existing difference in the state of cysteine residues between the two groups before sample treatment.

Sulfhydryl groups in cysteine residues form disulfide bonds through an oxidation reaction, which are essential for maintaining protein stability [16]. Serum total thiol, free thiol levels, disulfide/native thiol percentage ratios, and serum ischemia-modified albumin levels in patients with methamphetamine use disorders were significantly higher than those in the healthy control group. This suggests that these patients are under oxidative stress [17]. And oxidative stress plays an important role in methamphetamine-mediated neurotoxicity [18].

The number of modifications occurring in each differentially modified peptide was calculated, with detailed information provided in Table S3. Peptides with more than four modifications are shown in Table 5. Albumin was the most abundant protein in urine, accounting for 11%. And the abundance of protein AMBP in urine ranks third after albumin and uromodulin, accounting for 4% [19]. Due to the high abundance of these proteins in urine, it is likely that the peptides they contain were identified more frequently for modification. However, this study also found that, despite the low abundance of some proteins in urine, they still contained peptides that were modified a relatively large number of times.

**Table 5.**
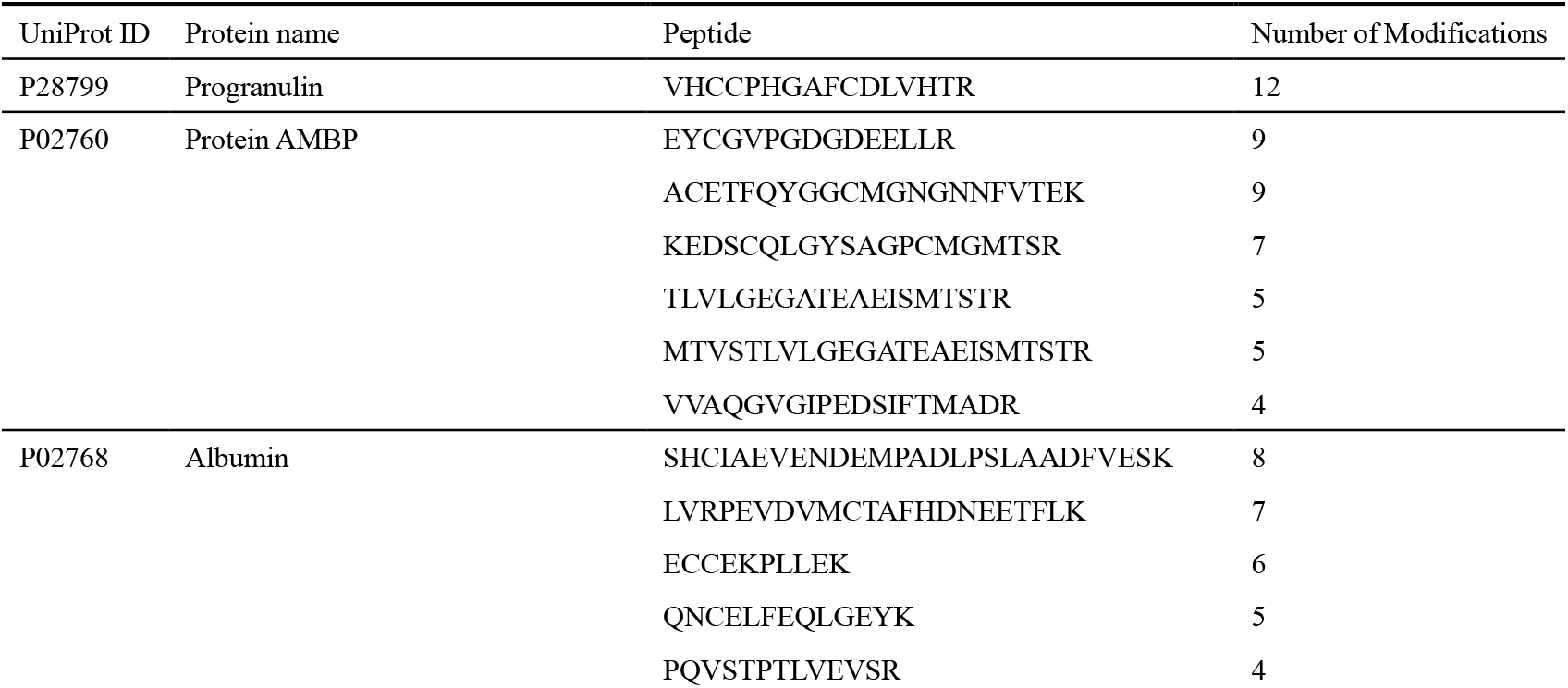
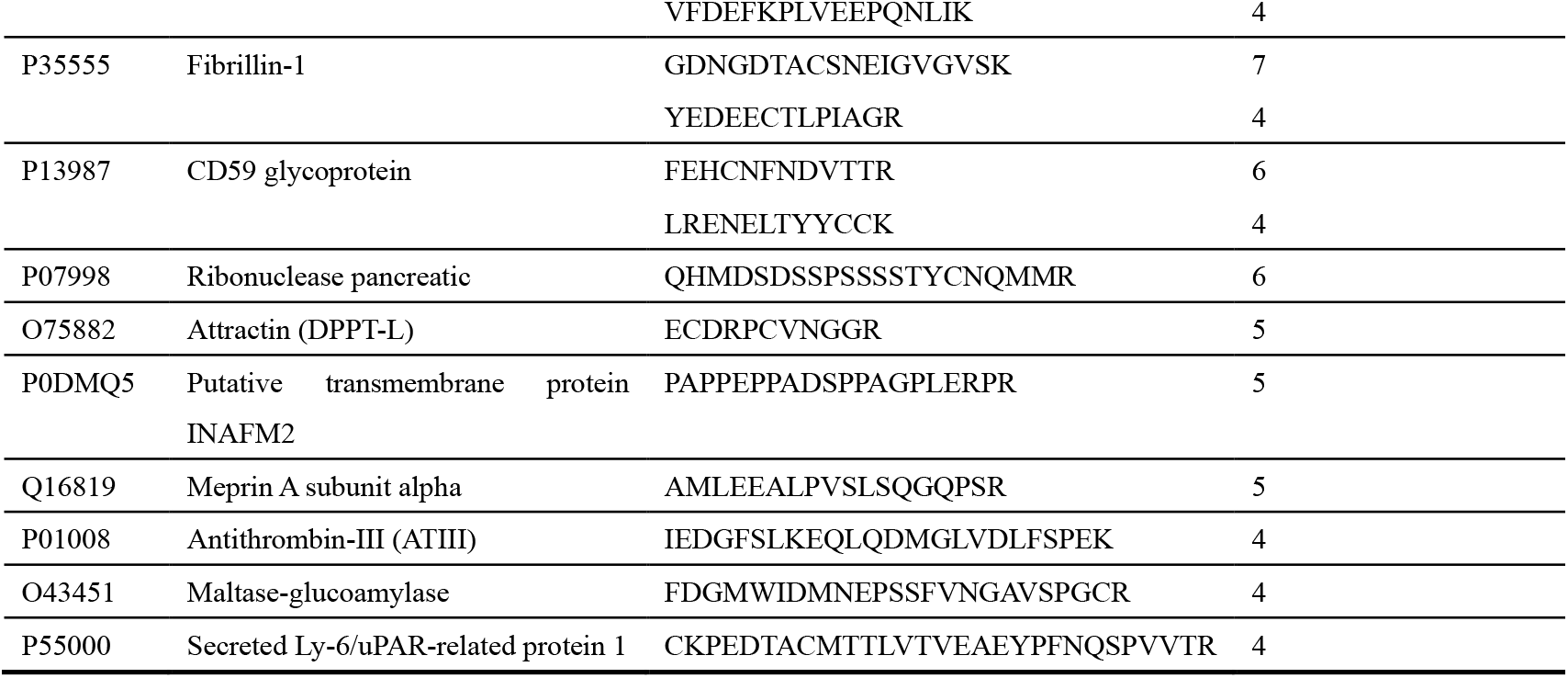
Differentially modified peptides with ≥ 4 modifications.

### 3.4. Analysis of Proteins Containing Differentially Modified Peptides

A total of 342 proteins containing differentially modified peptides were screened, with details provided in Table S4.

The number of species of differentially modified peptides identified in each protein was calculated, with detailed information provided in Table S5. Proteins containing ≥9 differentially modified peptides are shown in Table 6. Among them, albumin and protein AMBP contain more differentially modified peptides than other proteins. Similarly, the relatively high abundance of these two proteins in urine may be the reason for the relatively high number of identified differentially modified peptides they contain [19]. However, in this study, some proteins with lower abundance in urine were found to contain more differentially modified peptides than uromodulin, which ranks second in abundance. This group of lower-abundance proteins, yet with a high number of differentially modified peptide species, may reflect significant changes brought about by the modification of the urinary proteome in methamphetamine-abstinent rehabilitation patients. Some of these proteins have been reported to be associated with methamphetamine or amphetamine, while others that have not yet been reported to be associated with methamphetamine, may provide clues for the study of methamphetamine addiction rehabilitation.

**Table 6.**
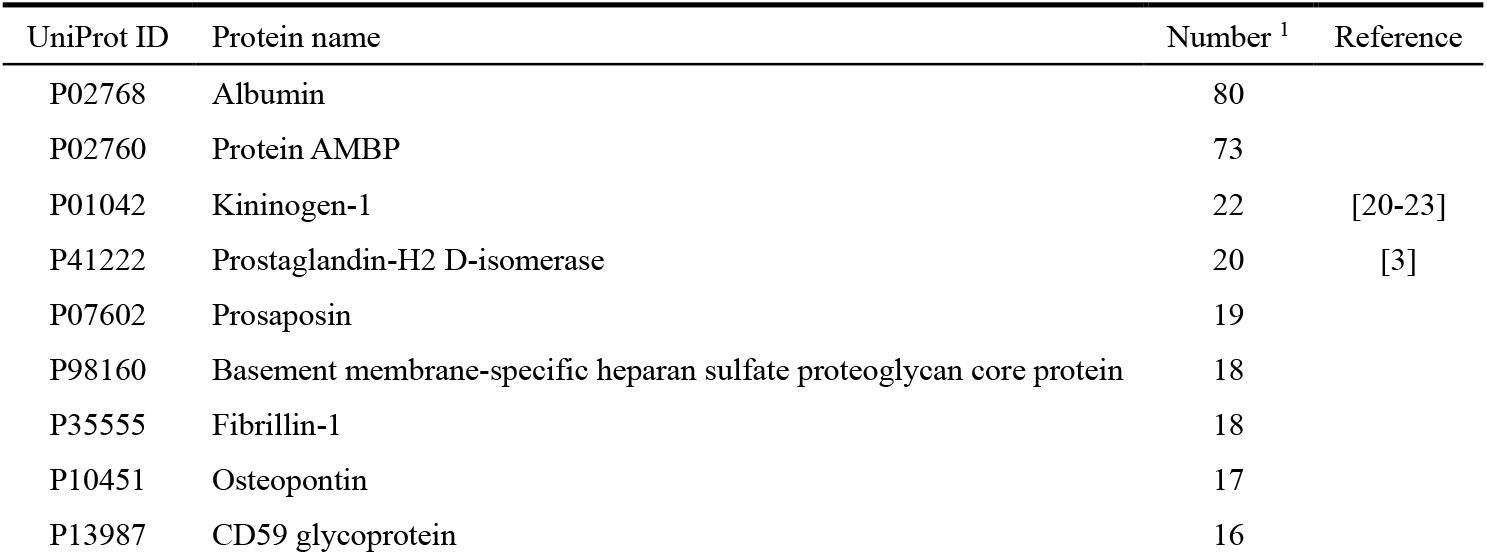
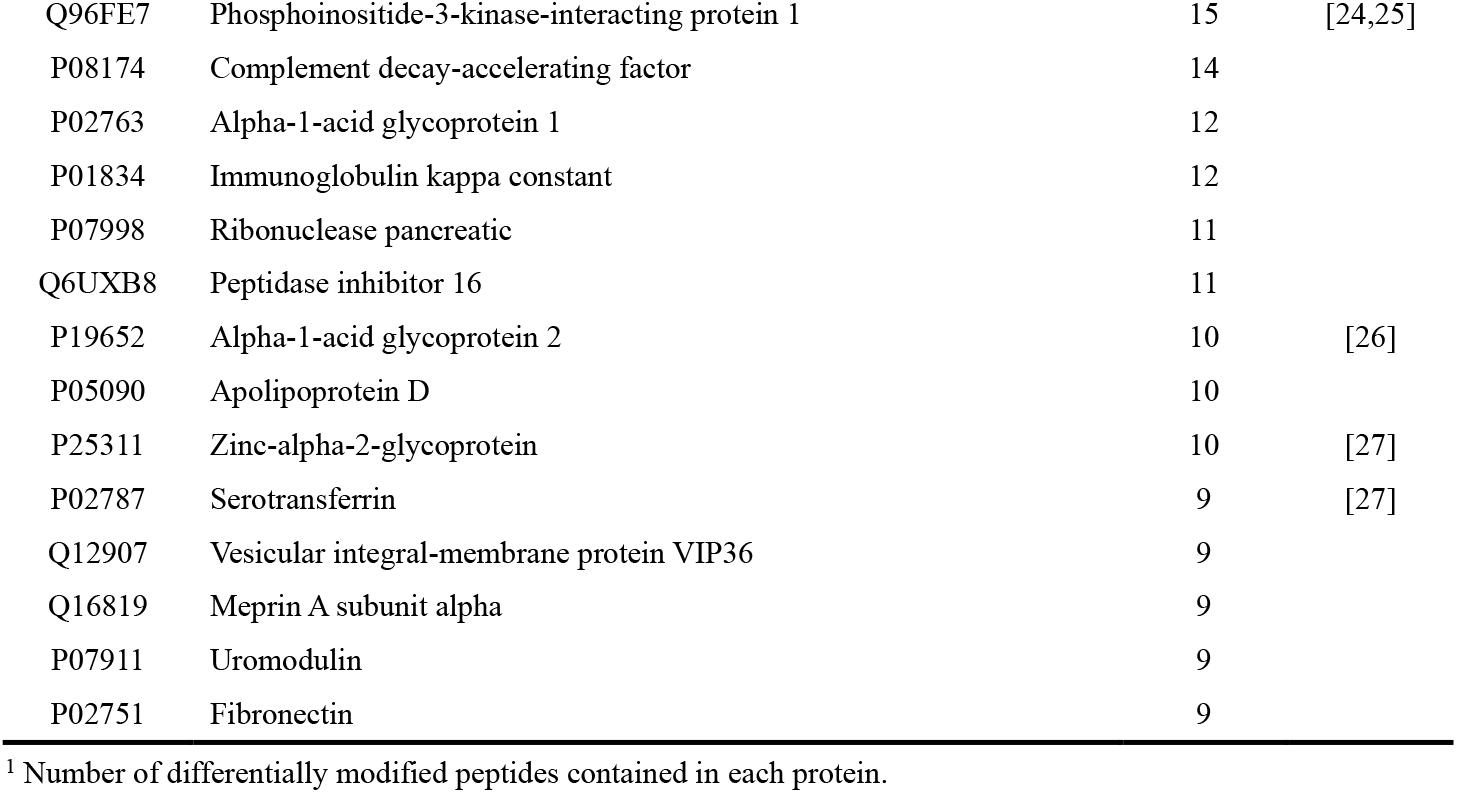
Proteins containing ≥ 9 types of differentially modified peptides.

Additionally, several proteins differ from those identified in our previous urinary proteome study of methamphetamine-abstinent rehabilitation patients (Table S6) [13]. The modifications that occurred in these proteins may reflect long-term changes in the body.

Functional analysis of the 342 proteins was performed using the UniProt website and the PubMed database. Several proteins have been reported to be associated with methamphetamine or amphetamine. Information of the significantly changed differentially modified peptides contained in these proteins is provided in Table 7, with additional details listed in Table S1.

**Table 7.**
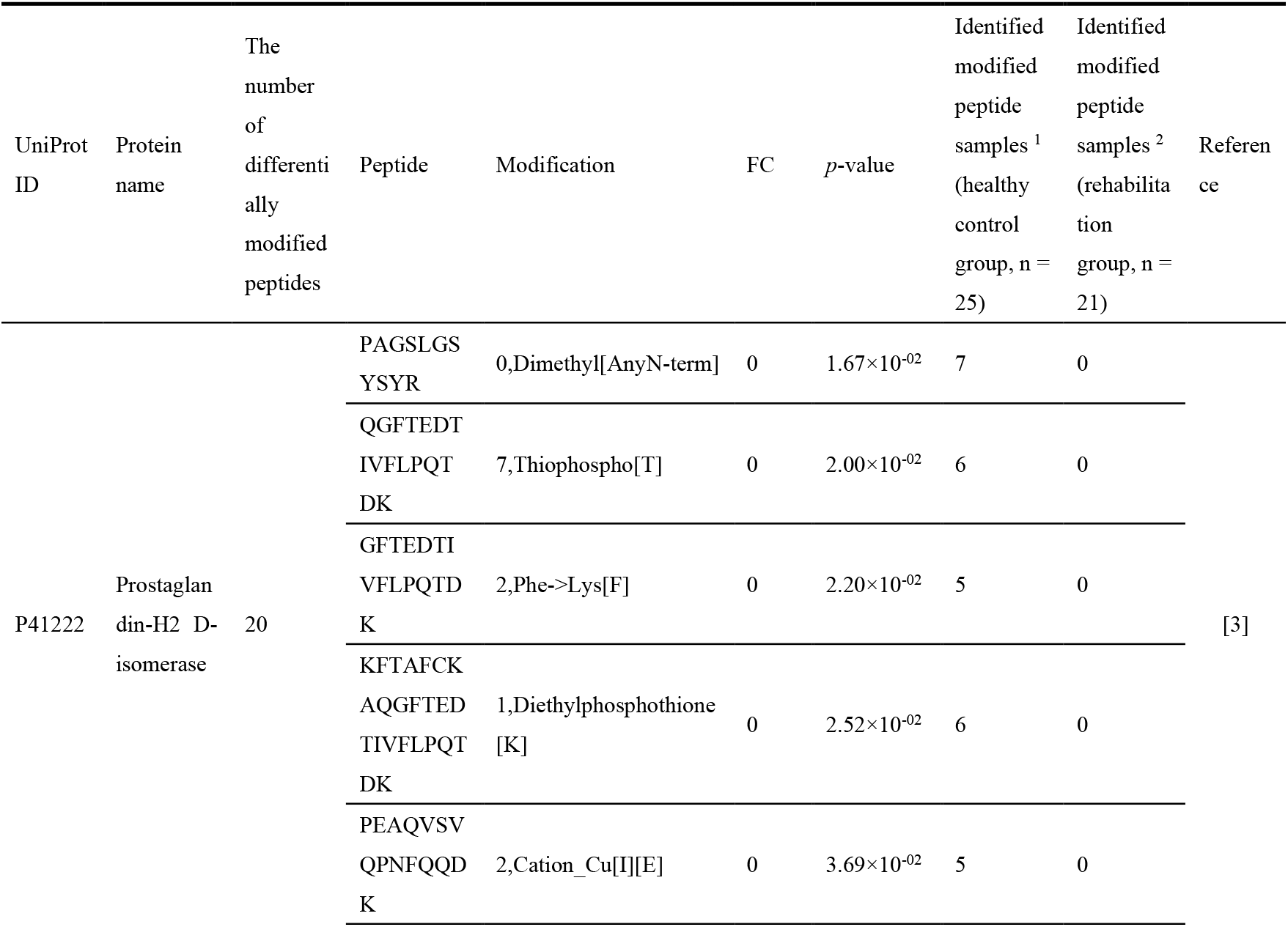
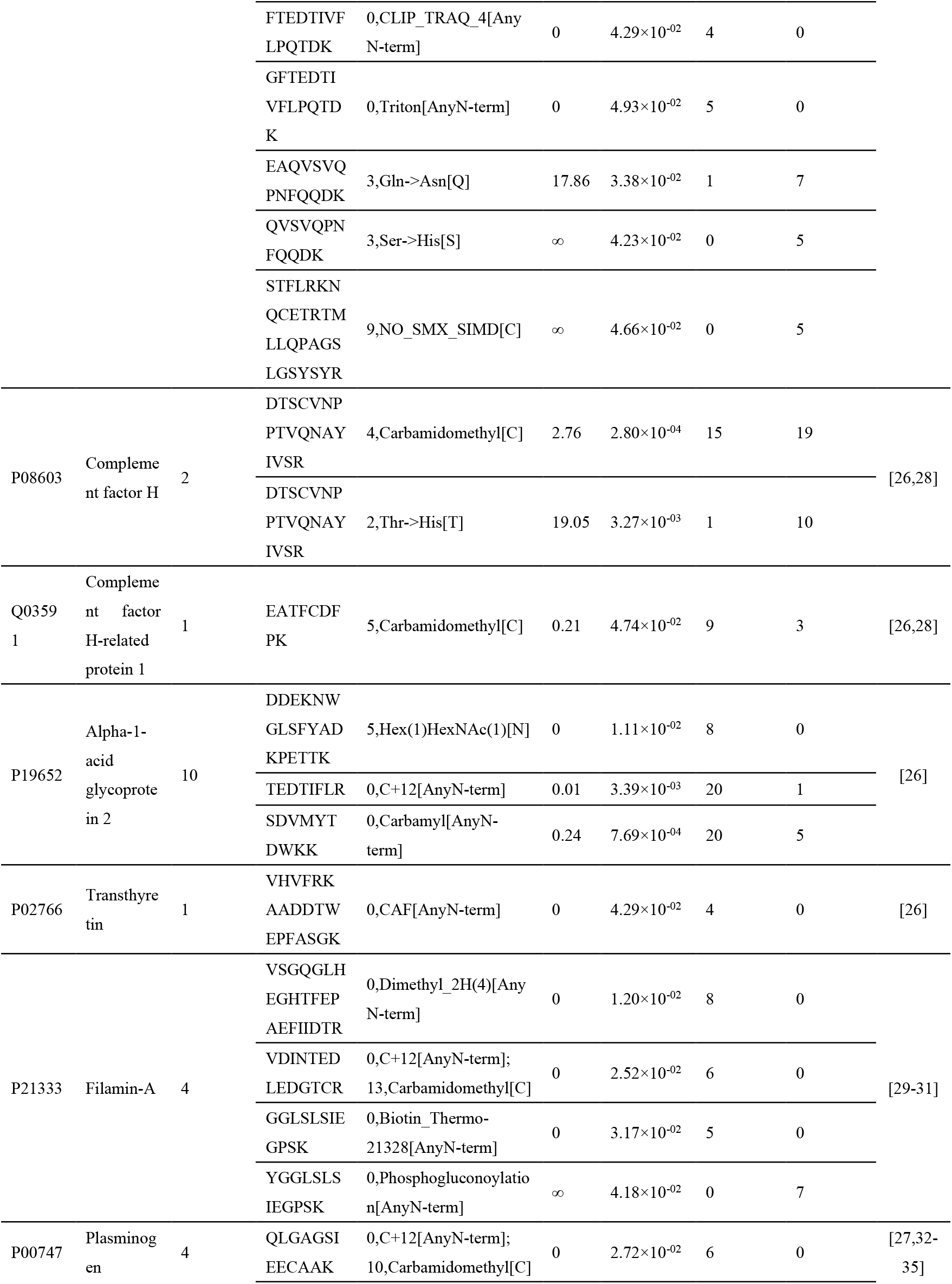
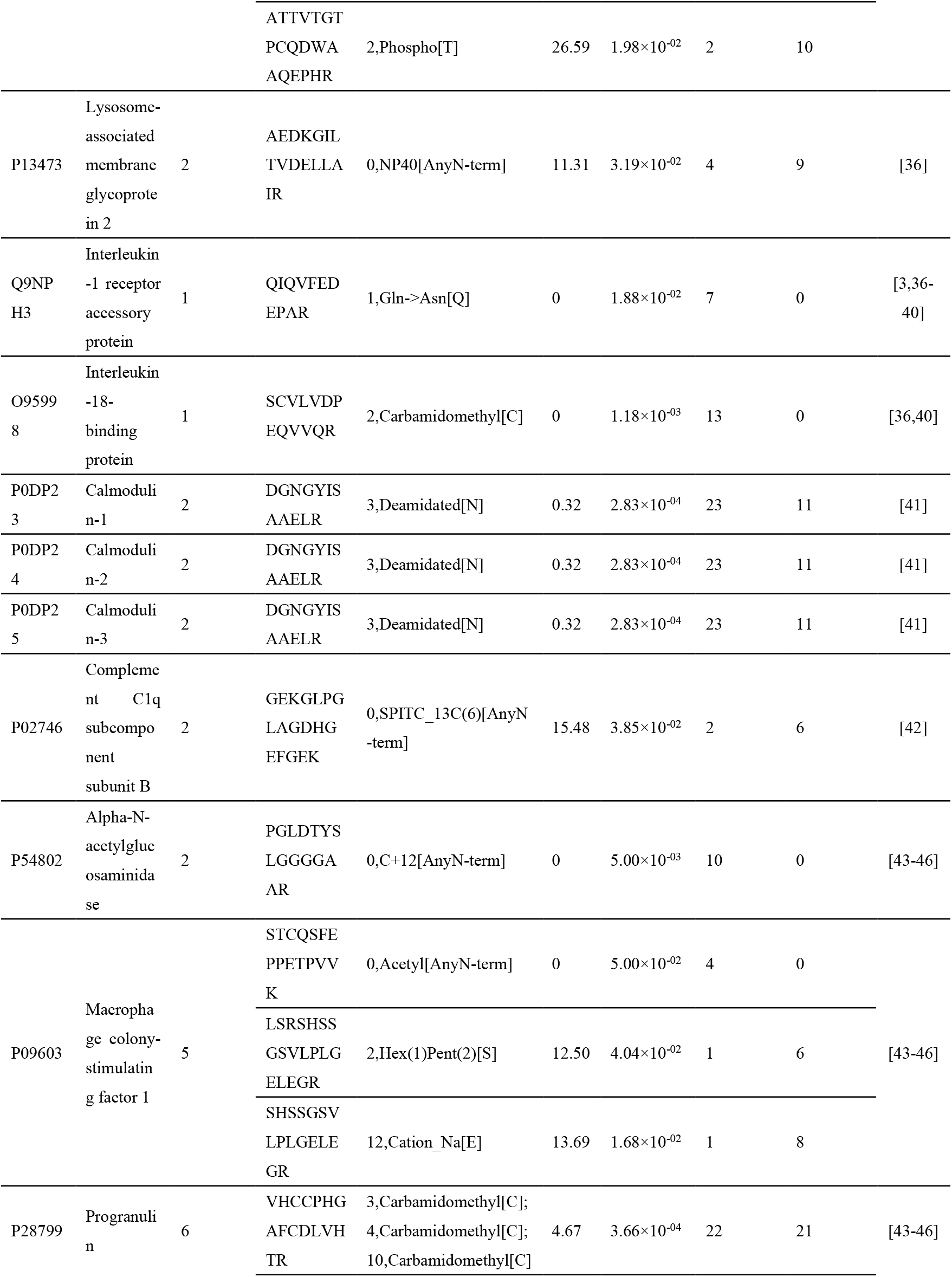
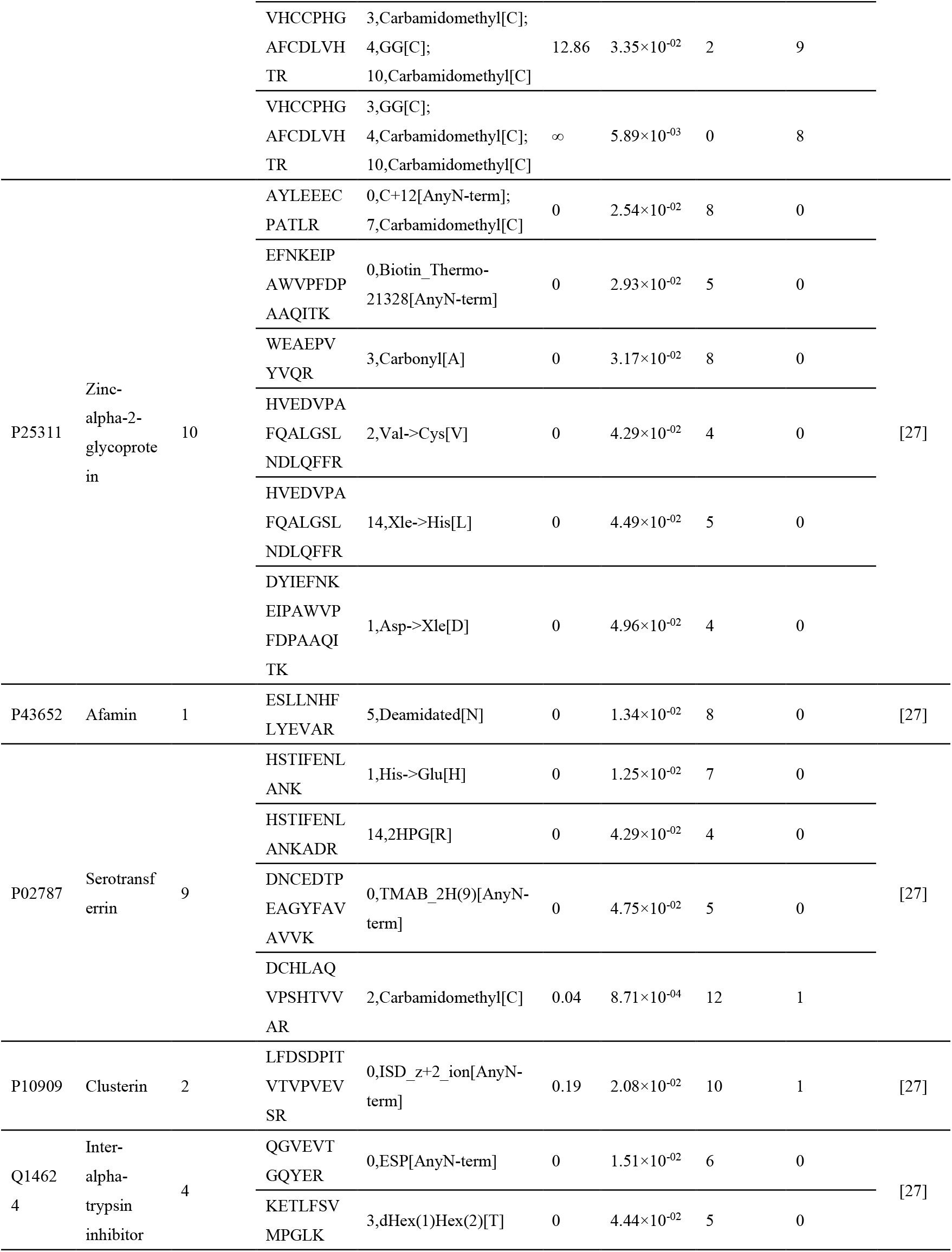
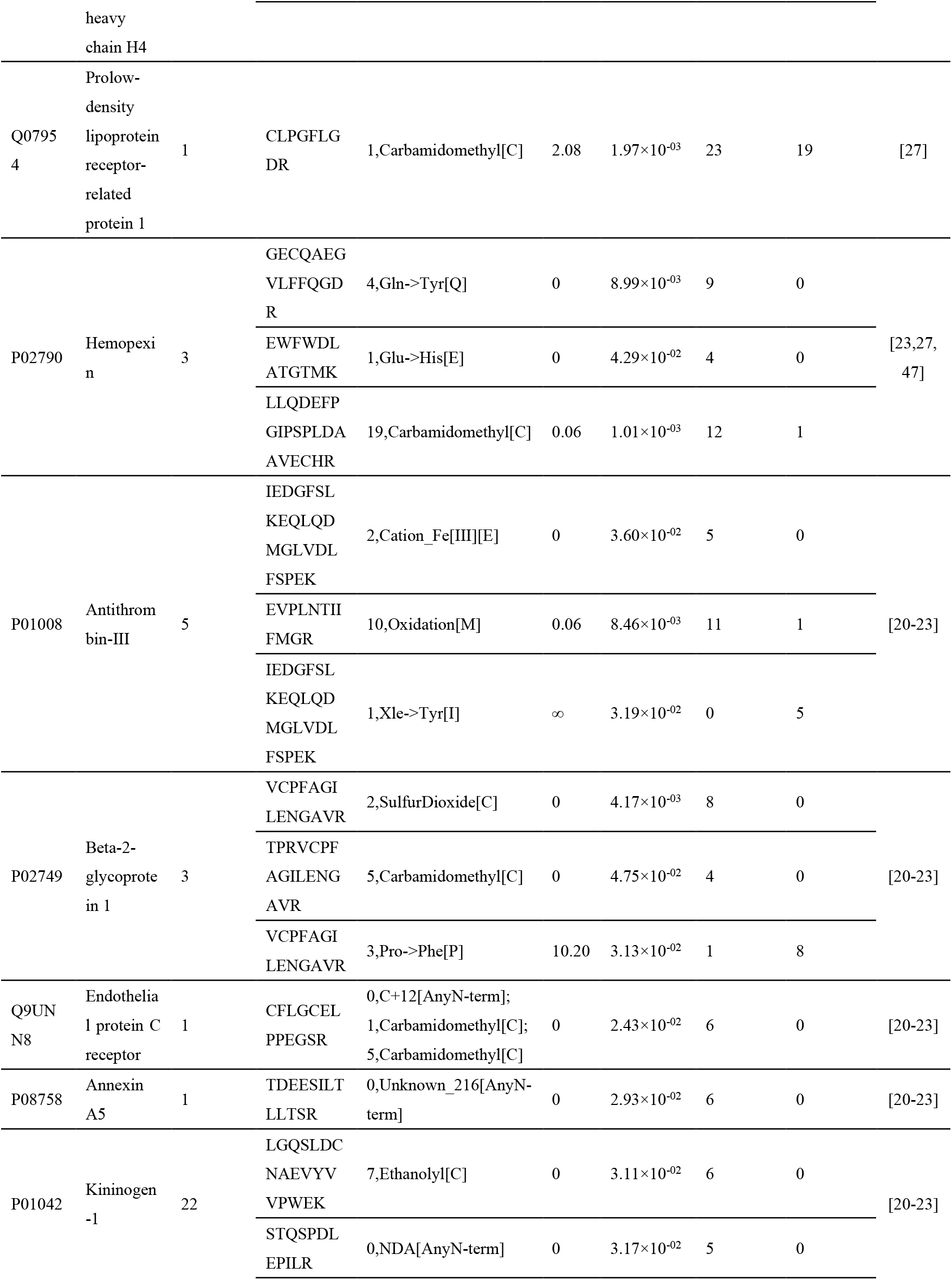
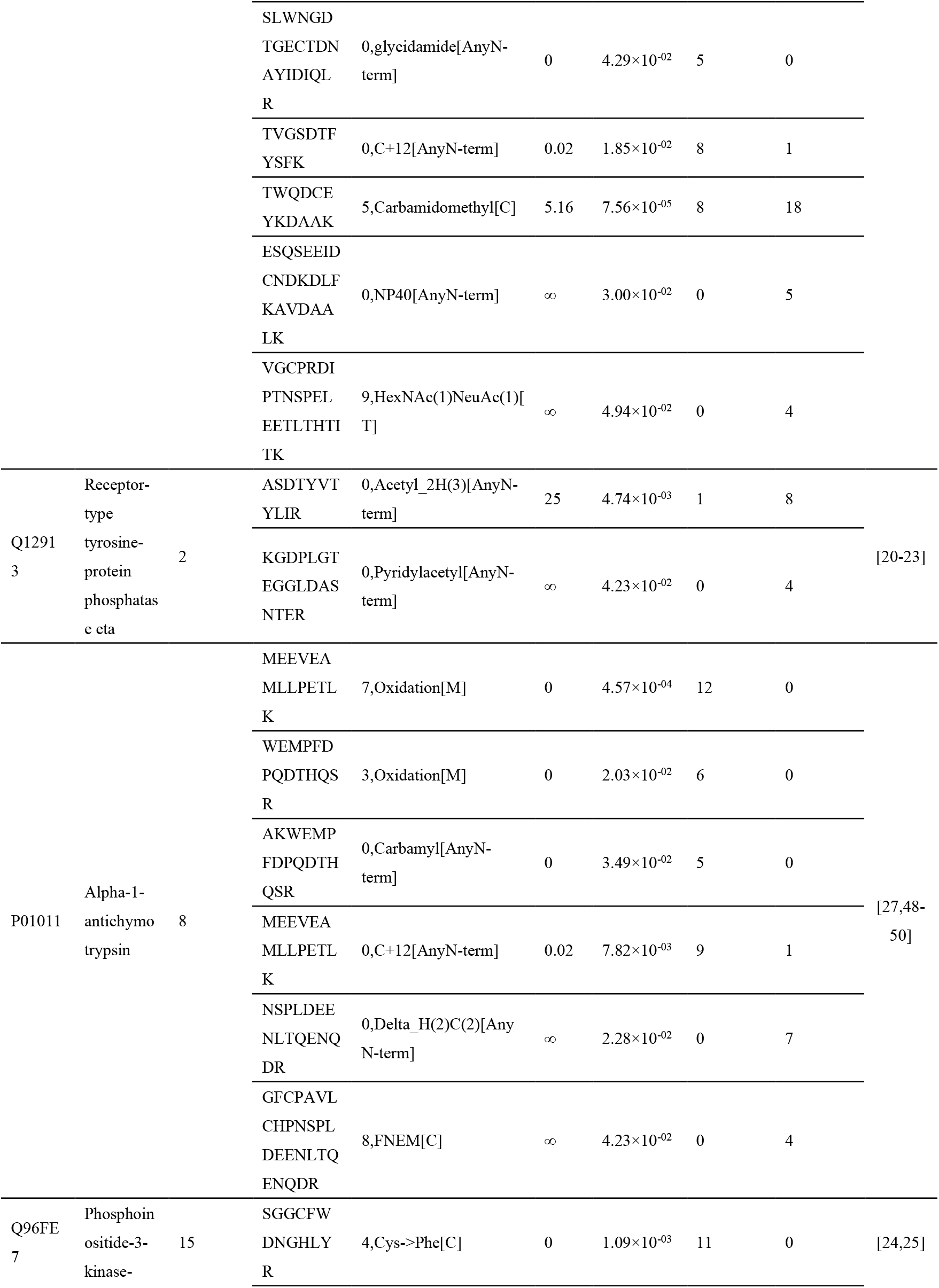
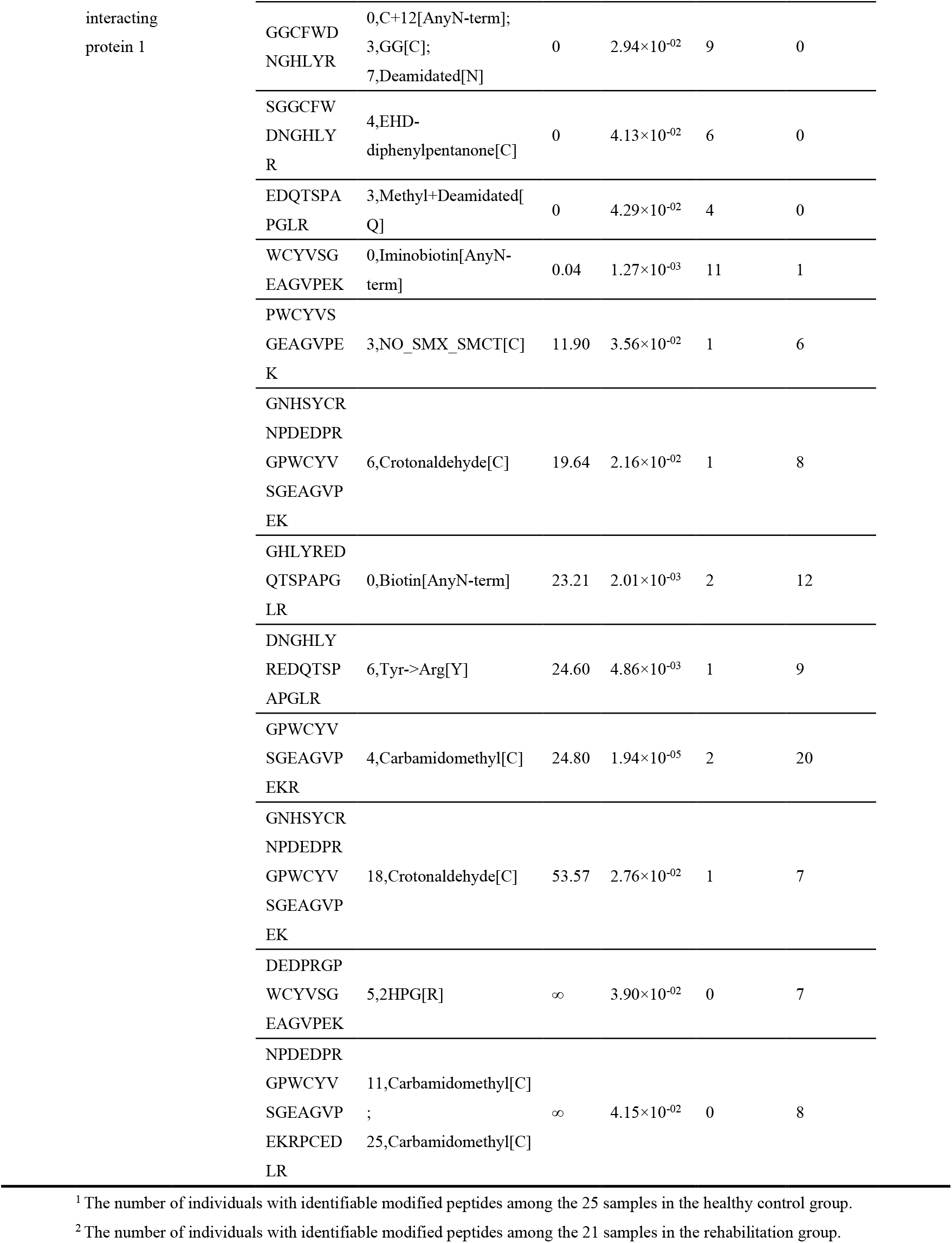
Information of significantly changed differentially modified peptides contained in proteins reported to be associated with methamphetamine or amphetamine.

#### (1) Prostaglandin-H2 D-isomerase

Methamphetamine withdrawal interferes with sleep-wake behavior and circadian rhythms, and sleep disturbances, in turn, increase methamphetamine use and the risk of addiction, creating a vicious cycle that heightens the risk of relapse [3]. Prostaglandin-H2 D-isomerase, which plays a role in regulating the circadian sleep/wake cycle, is low in abundance in urine but contains 20 identified differentially modified peptides. In contrast, uromodulin, which ranks second in abundance in urine, contains only 9 identified differentially modified peptides. This group of proteins, despite their low abundance, contains a high number of differentially modified peptide species, which may reflect significant changes brought about by urinary proteome modifications in methamphetamine-abstinent rehabilitation patients.

#### (2) Complement factor H, Complement factor H-related protein 1 (FHR-1), Alpha-1-acid glycoprotein 2 (AGP 2), Transthyretin

In the serum of methamphetamine addicts, the expression levels of complement factor H, AGP 2, and transthyretin are up-regulated. Complement factor H, a member of the complement activation family of regulators, is also up-regulated in the serum of methamphetamine-addicted rats, as well as in the rat hippocampus and ventral tegmental area. It may serve as a potential biomarker for methamphetamine addiction [26]. The up-regulation of complement factor H in chronic methamphetamine dependence may be associated with neuronal activity involved in addiction-related memory retrieval [28].

#### (3) Filamin-A, FLN-A

Filamin-A is involved in the adenylate cyclase-inhibiting dopamine receptor signaling pathway, which is initiated by the binding of dopamine to its receptor. Filamin-A specifically interacts with D2 and D3 dopamine receptors, influencing receptor localization and signaling, which may provide clues for the treatment of related neurological disorders [29].

In adenylyl cyclase (AC) 1/8-deficient mice, methamphetamine-induced behavioral sensitization was significantly attenuated, and sensitivity to methamphetamine in the nigrostriatal pathway was decreased, resulting in a weakened dopaminergic response. This indicates that AC 1/8 plays a role in methamphetamine-induced neuroplasticity and behavioral responses [30].

In addition, methamphetamine competes with dopamine for dopamine transporter (DAT) into dopaminergic terminals. It can lead to abnormal extracellular accumulation of dopamine by directly inhibiting DAT activity or reversing its direction. Methamphetamine also disrupts the proton gradient of vesicular monoamine transporter protein 2 (VMAT2) into synaptic vesicles, resulting in excessive dopamine release. It can also bind to monoamine oxidase (MAO) and block dopamine degradation, increasing the availability of free dopamine [31].

#### (4) Plasminogen

Plasminogen levels have been found to be elevated in the serum of patients with amphetamine addiction [27]. Plasminogen plays a role in trans-synaptic signaling by brain-derived neurotrophic factor (BDNF) and in modulating synaptic transmission. BDNF, which is abundant in the adult brain, is important for neuronal development, maintenance, transmission, and plasticity [32]. It has been suggested as a potential marker for methamphetamine addiction, with low levels of BDNF observed in many patients with addictive disorders, while elevated levels are seen in recovering methamphetamine addicts [33]. BDNF also influences the activity of the nucleus accumbens, inducing changes in dendritic spine density and triggering drug-related behaviors, such as self-administration and relapse [34]. Additionally, tissue plasminogen activator (tPA) mRNA expression in the frontal cortex, nucleus accumbens, striatum, and hippocampus increased in a dose-dependent manner after repeated methamphetamine treatments, and tPA activity in the nucleus accumbens was enhanced [35]. Moreover, tPA is a serine protease that activates plasminogen into fibrinolytic enzymes.

#### (5) Lysosome-associated membrane glycoprotein 2 (LAMP-2), Interleukin-1 receptor accessory protein, Interleukin-18-binding protein (IL-18BP)

Methamphetamine induces intestinal inflammatory injury through the overexpression of Nod-like receptor protein 3 (NLRP3) inflammasome, which regulates and directly activates IL-18 and IL-1β, playing an important role in both innate and adaptive inflammatory responses [36]. LAMP-2 is involved in the negative regulation of NLRP3 inflammasome complex assembly.

Another study showed that methamphetamine significantly increased IL-1β levels in the hippocampus of mice, while cognitive decline in methamphetamine-exposed mice could be attenuated by neutralizing IL-1 signaling. This suggests that cognitive decline associated with methamphetamine exposure may be correlated with IL-1 levels [37]. In addition, methamphetamine-induced relapse significantly increased the expression of IL-1β and TNF-α in the prefrontal cortex and hippocampus of rats. And cannabidiol treatment significantly reduced the expression of these pro-inflammatory cytokines, which may play an important role in inhibiting methamphetamine relapse [38]. Cannabidiol, a non-psychoactive component of cannabis with anti-inflammatory and immunosuppressive properties, reduces the motivation to self-administer methamphetamine and inhibits methamphetamine-induced relapse in rats [39].

Methamphetamine also stimulates microglia to release the cytokine IL-1β, a potent sleep regulator that exerts its effects through the interleukin 1 receptor [3].

In addition, serum levels of IL-18, TNF-α, and IL-6 were significantly elevated in previously chronic but currently abstinent methamphetamine users (mean number of days since last methamphetamine use was 39.06 ± 7.48 days). IL-6 levels in these users were significantly and positively correlated with Beck Depression Inventory (BDI) scores and the frequency of current methamphetamine use. Elevated levels of these pro-inflammatory cytokines may contribute to the psychopathological symptoms associated with chronic methamphetamine use [40]. And interleukin-1 receptor accessory protein is involved in the interleukin-1 mediated signaling pathway and the positive regulation of interleukin-6 production.

#### (6) Calmodulin-1, Calmodulin-2, and Calmodulin-3

Methamphetamine up-regulates calmodulin (CaM) expression and activates calmodulin-dependent protein kinase II (CaMKII), leading to neurological damage through the Ca^2^?-CaM-CaMKII signaling pathway [41].

#### (7) Complement C1q subcomponent subunit B

C1q is the initiating protein of the classical complement cascade, and its expression is significantly reduced in the hippocampal mossy fibers of methamphetamine-induced behaviorally sensitized mice. This reduction may be associated with the addiction process [42].

#### (8) Alpha-N-acetylglucosaminidase, Macrophage colony-stimulating factor 1 (CSF-1), Progranulin

Alpha-N-acetylglucosaminidase is involved in microglial cell activation. CSF-1 plays a role in microglial cell proliferation and the positive regulation of microglial cell migration. And progranulin is involved in the negative regulation of microglial cell activation.

Methamphetamine-induced neurotoxicity, accompanied by microglial cell activation, is characterized by proliferation, morphological changes, migration, and changes in inflammatory secretory activity [43]. The activation of microglia precedes pathological changes in dopaminergic axons in the striatum [44], and it persists for at least two years after withdrawal [45]. Microglial activation serves as a specific pharmacological marker of methamphetamine-induced neurotoxicity and is an important target for the treatment of methamphetamine addiction and related neural damage [46].

#### (9) Zinc-alpha-2-glycoprotein, Afamin, Serotransferrin, Clusterin, Inter-alpha-trypsin inhibitor heavy chain H4, Alpha-2-macroglobulin receptor

In the serum of amphetamine-addicted patients, several proteins were significantly up-regulated, including zinc-alpha-2-glycoprotein (up-regulated 3.49-fold, *p* = 0.007), afamin (up-regulated 2.84-fold, *p* = 0.004), serotransferrin (up-regulated 2.79-fold, *p* = 0.007), clusterin (up-regulated 2.74-fold, *p* = 0.037), ceruloplasmin (up-regulated 2.87-fold, *p* = 0.044), inter-alpha-trypsin inhibitor heavy chain h4 (up-regulated 1.6-fold, *p* = 0.053), and alpha-2-macroglobulin (up-regulated 2.77-fold, *p* = 0.01). Alpha-2-macroglobulin was found in more than one spot on the gels, which may be due to post-translational modifications, enzymatic effects, or the presence of different protein species [27].

#### (10) Hemopexin

Hemopexin was significantly up-regulated in the serum of amphetamine-addicted patients (up-regulated 3.17-fold, *p* = 0.008) [27]. Hemopexin interacts with macrophages to down-regulate lipopolysaccharide-induced production of tumor necrosis factor-α (TNF-α) and interleukin-6 (IL-6) [47]. Continued methamphetamine use in HIV-infected patients up-regulates hemopexin expression, which correlates with methamphetamine-induced oxidative stress [23].

#### (11) Antithrombin-III, Beta-2-glycoprotein 1, Endothelial protein C receptor, Annexin A5, Kininogen-1, Receptor-type tyrosine-protein phosphatase eta

Studies have shown that symptoms such as disseminated intravascular coagulation (DIC) have been observed in five patients who were intravenously administered phenmetrazine or methamphetamine [20], as well as in dogs that were intentionally fed methamphetamine [21]. DIC is a syndrome of systemic coagulation disorders triggered by various diseases, beginning with the activation of coagulation mechanisms [22]. Methamphetamine use also disrupts the coagulation pathway in HIV-infected individuals. In HIV-infected patients who continuously use methamphetamine, significant changes in the expression of antithrombin-III have been observed, and these changes cannot be attributed to HIV itself [23]. And antithrombin-III plays an important regulatory role in the coagulation system.

In addition, several proteins containing differentially modified peptides with significant changes identified in this study are involved in coagulation-related biological pathways. For example, beta-2-glycoprotein 1 is involved in the intrinsic pathway of blood coagulation and both the negative and positive regulation of coagulation. Endothelial protein C receptor, annexin A5 and kininogen-1 are all involved in blood coagulation and the negative regulation of coagulation. It has been shown that methamphetamine use triggers the up-regulation of kininogen in HIV-infected individuals [23]. And receptor-type tyrosine-protein phosphatase eta is involved in biological processes such as blood coagulation.

#### (12) Alpha-1-antichymotrypsin, ACT

Studies have found that ACT is up-regulated in the serum of patients with amphetamine addiction [27]. ACT is a serine protease inhibitor that co-localizes with Aβ in senile/neuritic plaques and is a potential inflammatory marker for Alzheimer’s disease [48]. It also serves as a diagnostic biomarker for amnestic mild cognitive impairment [49]. Methamphetamine increases the risk of developing Alzheimer’s disease through several pathways, such as strongly affecting the hippocampus-dependent regulation of reward circuits, inhibiting hippocampal neurogenesis, inducing long-term damage to hippocampal cells, and disrupting the structure and function of the blood-brain barrier. Additionally, addiction-associated mutations have been associated with an increased risk of neurodegenerative diseases, cognitive decline, and structural changes in the brain [50].

#### (13) Phosphoinositide-3-kinase-interacting protein 1

PI3K Interacting Protein 1 (PIK3IP1) acts as an inhibitor of the PI3K/Akt/mTOR pathway [24]. Methamphetamine inhibits the phosphorylation cascade of kinases in the PI3K-Akt-mTOR signaling pathway, leading to astrocyte apoptosis and autophagy. This pathway has also been demonstrated to be involved in addiction and cue-induced relapse related to several substances [25].

In this study, the differentially modified peptides associated with methamphetamine or exhibiting significant changes, along with the proteins in which they are located, provide insights into exploring drug addiction rehabilitation. However, this preliminary study focuses on comparisons of urinary proteome modifications. Further investigation is needed to understand how these modifications specifically regulate relevant biological processes at the cellular and molecular levels. Additionally, the sample size in this study is relatively small, and all the methamphetamine-abstinent rehabilitation patients included in the analysis were male. Further validation can be conducted through large-scale clinical studies in the future.

## 4. Conclusion

Significant differences in urinary proteome modifications were observed between drug rehabilitation patients who had abstained from methamphetamine for over three months and healthy individuals. These rehabilitation patients with over three months of methamphetamine abstinence still failed to return to the normal levels of healthy individuals, which may be used to shed light on the reasons for the high rate of methamphetamine relapse. Multiple proteins containing differentially modified peptides have been reported to be associated with methamphetamine. Additionally, several significantly changed modified peptides, not previously reported to be associated with methamphetamine, as well as the proteins they are located in, may provide new insights for research on the rehabilitation of methamphetamine addiction. The modifications that occur in these proteins may reflect long-term changes in the body.

This study presents a novel approach to exploring drug addiction through urinary proteome modifications, offering a new perspective on the study of drug withdrawal. It holds potential to provide clues for tracking and evaluating the drug addiction rehabilitation process.

